# Neural representation of emotional valence in human amygdala

**DOI:** 10.1101/2025.08.14.670371

**Authors:** Ke Bo, Lihan Cui, Gang Chen, Andreas Keil, Mingzhou Ding

**Affiliations:** Department of Psychological and Brain Sciences, Dartmouth College, Hanover, NH; J. Crayton Pruitt Family Department of Biomedical Engineering, University of Florida, Gainesville, FL; Department of Psychology, University of Florida, Gainesville, FL; Scientific and Statistical Computing Core, National Institute of Mental Health, Bethesda, MD

## Abstract

The amygdala is a core structure for encoding the affective value of external stimuli. Animal studies suggest that positive and negative emotions are separately encoded by distinct neuronal populations within the amygdala; however, this hypothesis has rarely been tested in humans. The current study examined this hypothesis by comparing the distributed emotion encoding model, as proposed in animal studies, with the univariate emotion encoding model using functional magnetic resonance (fMRI) imaging. More specifically, we applied univariate regression, using average amygdala activation to represent global activation level, and multivariate regression, using distributed voxel-level pattern within the amygdala, to predict normative valence of affective images from the IAPS library. In the core amygdala, the multivariate model’s prediction performance was not better than that of the univariate model, with weight map analysis revealing an overwhelming predominance of voxels selectively responsive to negative stimuli. When the region of interest was expanded to include voxels with lower anatomical probability of belonging to the amygdala as well as voxels from adjacent areas, the multivariate model significantly outperformed the univariate model, with the voxels selectively responsive to positive valence primarily located in regions surrounding the core amygdala. These findings suggest that in the human amygdala, the core region encodes emotional valence primarily through a global activation signal, rather than distributed patterns consisting of separate clusters of positive and negative voxels, and a more distributed valence representation emerges when voxels surrounding the amygdala are taken into consideration.

## Introduction

The amygdala is a central structure in the brain’s affective network. The types of emotions that maximally engage the amygdala, however, is debated. Early studies primarily focused on the amygdala as a hub for processing negative emotions, particularly fear (Breiter et al., 1996; Morris et al., 1996; Phan et al., 2002), and on its role in coordinating the rapid subcortical responses crucial for survival in threatening situations (Garvert et al., 2014; Méndez-Bértolo et al., 2016; Pessoa & Adolphs, 2010). Recent research has begun to expand this perspective, demonstrating that the amygdala is equally critical for encoding positive emotions. For instance, a large-scale meta-analysis of human neuroimaging data implicated the amygdala in various positive emotional states, such as happiness, humor, and sexual arousal (Costafreda et al., 2008). Additionally, more recent studies have underscored its role in reward learning (Aquino et al., 2020; Wassum, 2022) and food craving (Ghobadi-Azbari et al., 2023; Sun et al., 2015). How the polarity of emotion, often referred to as “valence,” is represented in the amygdala remains an active research question.

Emotional stimuli (positive as well as negative) are more intense than neutral stimuli. Univariate neuroimaging methods, which tend to treat the amygdala as an unitary functional structure and derive a single response measure, find that the overall amygdala activation encodes the intensity of emotion regardless of valence (Bonnet et al., 2015). Evidence from animal studies, however, suggests a more nuanced role for the amygdala in emotion encoding. A framework derived from the rodent model proposes that valence is represented in the amygdala through population coding (Headley et al., 2019). Specifically, there are distinct neural ensembles in the amygdala that respond preferentially to positive or negative emotions and the weighted combination of activities in these neural ensembles encodes the affective states with different valences. Rodent studies have further shown that positive- and negative-selective neurons rarely overlap (Corder et al., 2019; Gründemann et al., 2019; Kyriazi et al., 2018; Zhang & Li, 2018) and they connect to distinct downstream brain regions to facilitate the elaboration of differently-valanced inputs and enable adaptive responses (Oneill et al., 2018). To what extent a similar population coding scheme exists in the human amygdala is unclear. Human emotions are notably more complex, often involving intricate processes of appraisal and evaluation mediated by cortical regions (Lindquist & Barrett, 2012). It is conceivable that humans may represent emotional valence differently from those observed in animal models. Testing these has been challenging, not only because human studies primarily rely on functional MRI (fMRI), which captures activity at a coarser millimeter-scale resolution, but also because the prevailing univariate analysis of fMRI data has inherent limitations in revealing the more nuanced picture of emotion processing in the amygdala.

An emerging method for investigating population coding in humans is the multivariate pattern analysis (MVPA) (Kragel et al., 2018; Norman et al., 2006). It is typically done by using fMRI voxels as features to train classifiers or regression models to distinguish between experimental conditions or predict an outcome variable (e.g., valence of the stimulus). This technique treats voxels as functional units and takes into account of the multivoxel spatial pattern formed by voxel-level activations. Additionally, this approach enables the generation of weight maps, which allow us to examine whether specific voxels preferentially encode positive or negative emotions (Haufe et al., 2014), potentially bridging the gap between human fMRI and animal single unit studies. In this work, we propose to apply MVPA to fMRI data to examine the following contrasting models of valence representation in the amygdala: (1) If valence is encoded in the amygdala through population coding, we would expect that the multivariate analysis taking into account pattern information would significantly better predict emotional valence than the univariate analysis and (2) if the multivariate model fails to significantly better predict emotional valence, this would suggest that valence is represented by the overall activity level rather than by the distributed activity pattern. The weight map analysis can corroborate the conclusions by revealing the prevalence of voxels selective for different emotions: (1) if the population coding model applies, then we would expect to see a mixture of voxels selective for either negative or positive emotions and (2) if that is not the case, then the weight map would be dominated by voxels preferentially encoding only one type of emotion.

A recent study has attempted to use MVPA to classify emotions based on the response patterns in the amygdala but was unsuccessful (Varkevisser et al., 2023). A potential reason for this failure could be that the study restricted its analysis to the high-probability voxels from the probability atlas of the amygdala (Amunts et al., 2005), the so-called “core” amygdala, which may have limited the spatial variance in the neural patterns. In fact, meta-analyses of neuroimaging studies consistently report emotional activations not only in the core amygdala but also in its surrounding regions, including edge amygdala regions next to the hippocampus and parahippocampal areas (Sergerie et al., 2008). For example, one meta-analysis found that only 50% of peak fMRI activations elicited by emotional stimuli were located within the core amygdala, with the remaining activation peaks distributed along the edges of the amygdala and even regions outside of the amygdala (Ball et al., 2009). These findings suggest that functionally relevant emotional activations may exist beyond the core amygdala and should be taken into consideration when examining amygdala encoding of emotional information.

In this study, we aim to test to what extent valence is represented via a population (multivariate) model or a global activity (univariate) model, within the core amygdala as well as in the various expanded definitions of the amygdala. We recorded fMRI data while twenty participants passively viewed 60 images from the International Affective Picture System (IAPS; Lang et al., 1997). These images span the full spectrum of valence, with arousal levels carefully matched between the positive and negative emotions to minimize the potential confounding influence. Our first analysis examined population coding of valence within the core amygdala area by training a multivariate regression model to predict normative ratings of emotional valence for the IAPS images based on voxel-level activity patterns. To evaluate whether the multivariate prediction performance is driven by the overall amygdala activation, we compared this model’s performance to that of a univariate prediction framework using the average amygdala activation as the predictor. We then extracted the weight map from the multivariate analysis in the second analysis to identify voxel-level valence-specific coding within the amygdala, assessing the preferential coding of individual voxels for positive or negative valence. Third, we applied the above analyses to less constrained definitions of the amygdala by including voxels with smaller probability in the probabilistic atlas to explore whether distributed neural patterns or global activity levels encode valence within more broadly defined amygdala regions.

## Method

### Participants

The experimental protocol was approved by the Institutional Review Board of the University of Florida. Twenty-six healthy volunteers with normal or corrected-to normal-vision gave informed consent and participated in this experiment. Before the MRI scan, two subjects withdraw from the experiment. In addition, data from four participants were excluded due to excessive head movements in the scanner. The data from the remaining twenty subjects were analyzed and reported here (10 men and 10 women; mean age: 20.4±3.1).

### Stimuli

The stimuli consisted of 20 pleasant, 20 neutral and 20 unpleasant pictures selected from the International Affective Picture System (IAPS) (Lang et al., 1997); their IDs are: 20 pleasant pictures: 4311, 4599, 4610, 4624, 4626, 4641, 4658, 4680, 4694, 4695, 2057, 2332, 2345, 8186, 8250, 2655, 4597, 4668, 4693, 8030; 20 neutral pictures: 2398, 2032, 2036, 2037, 2102, 2191, 2305, 2374, 2377, 2411, 2499, 2635, 2347, 5600, 5700, 5781, 5814, 5900, 8034, 2387; and 20 unpleasant pictures: 1114, 1120, 1205, 1220, 1271, 1300, 1302, 1931, 3030, 3051, 3150, 6230, 6550, 9008, 9181, 9253, 9420, 9571, 3000, 3069. These pictures are categorized based on their normative valence and arousal levels (Liu et al., 2012, Bo et al., 2021, Bo et al., 2022). The pleasant pictures included sport scenes, romance, and erotic couples, whereas the unpleasant pictures included threat, attack scenes, and bodily mutilations. The neutral pictures included landscapes and neutral human beings. Across emotional contents, pictures were matched for presence/absence of living/nonliving content, as well as for landscape/scene versus close-up shots, and they were also matched for in-house ratings of perceived complexity obtained from several hundreds of undergraduate students. The normative arousal rating for unpleasant pictures (6.2 ± 0.79) was matched to that of pleasant pictures (5.8 ± 0.90), as reported in Bo et al. (2021).

### Experimental paradigm

See Figure 1A. In each trial, the IAPS picture was displayed on a MR-compatible monitor for 3s, which was followed by a variable interstimulus interval (2800 ms or 4300 ms). A scanning session was comprised of 60 trials corresponding to 60 different pictures. There were five sessions and the order of the same 60 pictures was randomized from session to session. Pictures, viewed via a reflective mirror, were presented on the monitor that was placed outside the scanner. Participants were instructed to maintain fixation on the center of the screen during the whole session. After the experiment, participants rated the hedonic valence and emotional arousal levels of 12 representative pictures (4 pictures from each of the three broad emotion categories) which were not displayed during the experiment. The rating was done using a paper and pencil version of the self-assessment manikin (Bradley and Lang, 1994).

**Figure 1.**
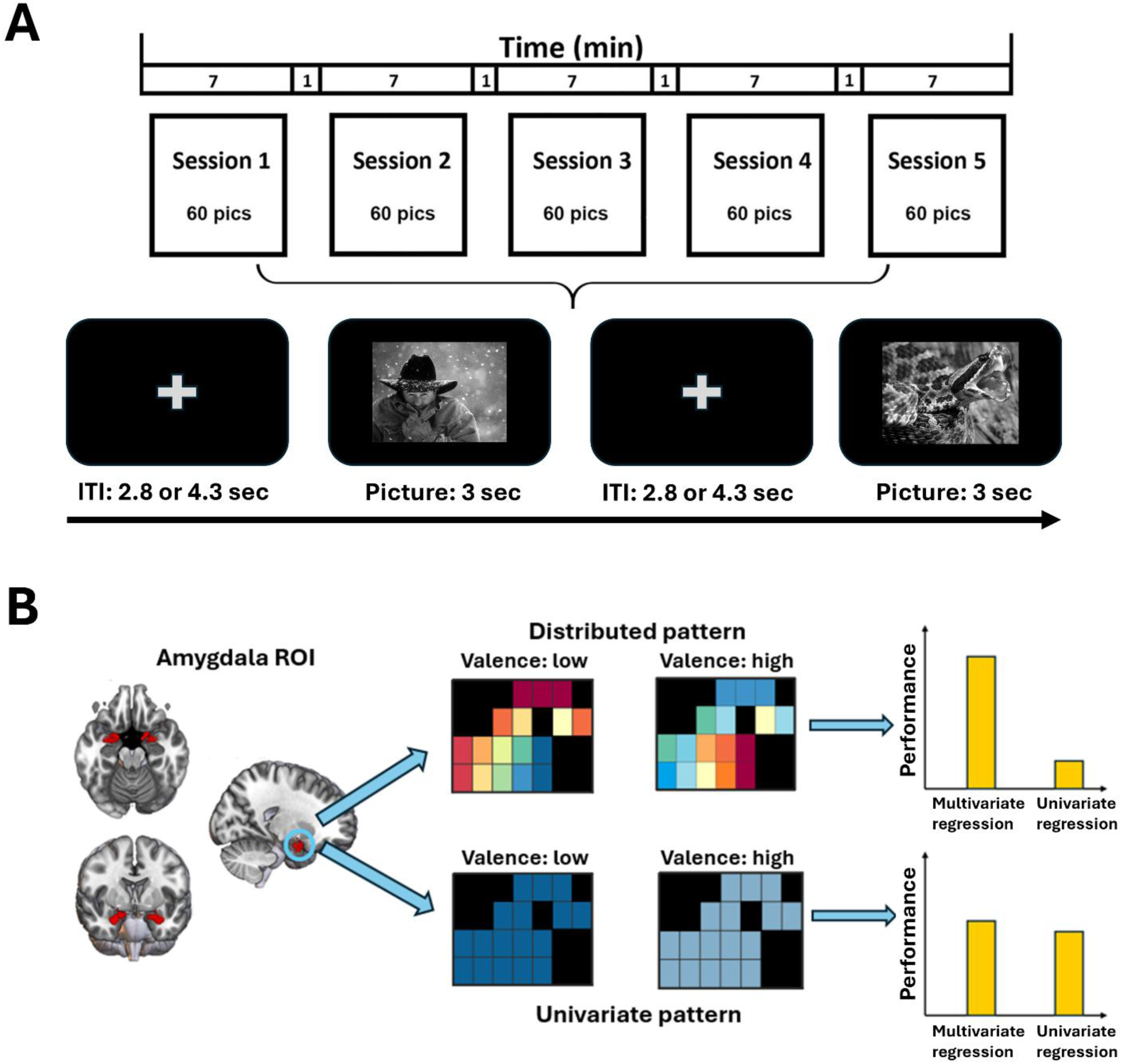
Experimental paradigm and analytical scheme for hypothesis testing. **(A)** There were five sessions. Each session lasted 7 min. A total of 60 IAPS pictures, including 20 pleasant, 20 unpleasant, and 20 neutral, were presented in each session, the order of which was random and also randomly varied from session to session. Each picture lasted 3 s and was followed by a fixation period referred to as intertrial interval (ITI) (2.8 or 4.3 s). **(B)** The distributed pattern hypothesis predicts that the multivariate regression model outperforms the univariate regression model in predicting normative valence. The univariate pattern hypothesis predicts that the multivariate regression model and the univariate regression model perform similarly in predicting normative valence.

### fMRI data acquisition

Functional MRI data were collected on a 3T Philips Achieva scanner (Philips Medical Systems), with the following parameters: echo time (TE), 30 ms; repetition time (TR), 1.98 s; flip angle, 80°; slice number, 36; field of view, 224 mm; voxel size, 3.5*3.5*3.5 mm; matrix size, 64*64. Slices were acquired in ascending order and oriented parallel to the plane connecting the anterior and posterior commissure. T1-weighted high-resolution structural images were also obtained.

### Data preprocessing

Preprocessing was done using SPM. The first five volumes from each session were discarded to eliminate artifacts caused by the transient instability of the scanner. Slice timing was corrected using interpolation to account for differences in slice acquisition time. The images were then corrected for head movement by spatially realigning them to the sixth image of each session, normalized and registered to the Montreal Neurological Institute (MNI) template, and resampled to a spatial resolution of 3 * 3 * 3 mm. The transformed images were smoothed by a Gaussian filter with a full width at half maximum of 8 mm. Other smoothing parameters were also applied to test the robustness of the findings. The low frequency temporal drifts were removed from the functional images by applying a high-pass filter with a cutoff frequency of 1/128 Hz.

### Beta-series estimation

BOLD activation was estimated on a trial-by-trial basis by using the beta series method (Mumford et al., 2012). In this method, a given trial was represented by a regressor, and all of the other trials were represented by another regressor. Six motion regressors were also included to account for any movement-related artifacts during the scan. Repeating the process for all the trials we obtained the BOLD response to each picture presentation in each of the brain voxels.

### Region of Interest (ROI)

Our main focus in this study is how the human amygdala represents valence. The amygdala ROI is defined according to a probabilistic map of the amygdala (Amunts et al., 2005; Mumford et al., 2012). Specifically, voxels within the centromedial, basolateral, and superficial subdivisions of the amygdala that have a probability greater than 50% were aggregated to construct the core amygdala ROI. In addition to the core amygdala, we also considered extended amygdala by progressively including voxels with probability thresholds of 30%, 10%, and 0%, as well as voxels adjacent to the 0% probability amygdala boundary.

### Linear regression analysis of picture-evoked amygdala responses

To investigate whether emotional valence was represented in a more distributed versus a more univariate fashion in the amygdala, linear multivariate as well as univariate regression analysis was conducted on the ROI-level and voxel-level activities in the core as well as the extended amygdala ROIs. For each participant, single-trial beta values from the five repetitions of the same IAPS picture were averaged to enhance signal-to-noise ratio, resulting in 60 picture-evoked responses per participant. The voxel-level activities and voxel-averaged univariate activation of the amygdala were employed as independent variables in separate regression models. The regression model was trained to predict the normative valence rating of each IAPS picture. The method combined Least Absolute Shrinkage and Selection Operator (LASSO) with leave-one-subject-out cross-validation. Specifically, the model was trained on the data from nineteen subjects and tested on the remaining one subject, and the process was repeated 20 times. The model performance was computed by averaging the 20 test performances. Since the analysis was between-subject, the data was normalized within each subject before fitting regression models to ensure that the contribution from each subject was at a similar level. The final regression model at the population level was established by averaging the 20 training models from the leave-one-subject-out cross-validation.

To assess the statistical significance of the model’s predictive performance, we conducted a permutation test. The correspondence between amygdala activity evoked by each stimulus and valence ratings was randomly shuffled. Repeating this procedure 1000 times, we obtained an empirical distribution of regression performance. The threshold for statistical significance was defined as the 95th percentile of this distribution (the top 50 values), corresponding to p = 0.05. The performance of the multivariate model (using all voxels within the ROI as predictors) and that of the univariate model (using voxel-averaged activity for the ROI as the predictor) were compared to assess whether a more distributed representation or a more univariate representation provided a better explanation of the data.

### Hypothesis testing and weight map analysis

The distributed-pattern and univariate-pattern hypotheses make distinct predictions with respect to the regression analysis. According to the population-coding model, if emotional valence is encoded by distributed spatial patterns, the performance of the multivariate model should exceed that of the univariate model in predicting normative valence. This prediction rests on the assumption that voxels are emotion-selective and exhibit differential responses to differently valenced input, yielding distinctive neural patterns for different emotions. Averaging across voxels—as in univariate modeling—diminishes the distinction between inputs of differing valences and thereby reducing model performance. In contrast, the univariate-pattern hypothesis predicts comparable regression performance for both the multivariate and the univariate models. This hypothesis assumes that all voxels share similar emotion selectivity, so that differences in average response magnitude suffice to encode valence and treating the amygdala response as a spatial pattern provides no additional information (Figure 1B).

From the foregoing, it is clear that to evaluate our hypotheses, it is important to assess the emotional selectivity of voxels in the amygdala. We achieved this by following the methodology proposed by Haufe et al. (2014). In a multiple regression model, a weight is assigned to every predictor variable in the form of the beta coefficient for the variable (e.g., activation of a voxel). It has been shown that the direct physiological interpretation of these weights is not straightforward. Voxels that do not contain task-related signals can be assigned high weights, while voxels containing task-related signals may be assigned low weights. To address this, Haufe et al. (2014) proposed a transformation that transforms the weight vector from the multiple regression model into a new vector according to, A = Cov(X) * W, where the X is the recorded data, W is the weight vector of the regression model, and A represents the transformed weight vector. After transformation, the sign of the weight reflects the emotion selectivity of the voxel, the weight magnitude reflects the strength of that selectivity. In the present experiment, if the valence is ordered from low (unpleasant) to high (pleasant), or 1 to 9, a positive weight indicates that the BOLD activation of the voxel is higher for higher valenced inputs, namely, it is selective for the positive emotion. Likewise, a negative weight indicates that the BOLD activation of the voxel is higher for lower valenced inputs, namely, it is selective for the negative emotion.

## Result

Participants viewed the presentation of 60 IAPS pictures from three broad categories: pleasant (20), neutral (20) and unpleasant (20) while their fMRI data were recorded. Each picture was associated with a normative valence. The goal of this study was to examine whether the normative valence was represented in a more distributed vs a more univariate fashion in the core amygdala and its adjacent structures.

### Core amygdala

The core amygdala is defined as the voxels with a greater than 50% probability of belonging to any amygdala subregion in the established probabilistic amygdala atlas (Amunts et al., 2005). To test models of valence coding within this ROI, we trained a multivariate pattern regression model using all voxels within the core amygdala as input features (predictor variables) and compared its performance to a univariate regression model that used the average activity within the core amygdala as the input feature (predictor variable). Here performance is defined to be the correlation between the predicted valence and the actual normative valence of the IAPS pictures. This comparison was aimed to assess: (1) whether normative valence can be predicted by the neural activity in the amygdala, and (2) which model yielded better predictions. Specifically, as shown in Figure 1B, assuming that both models can predict valence above chance level, (1) if the multivariate regression model’s performance is significantly better than the univariate regression model, this would suggest that the amygdala encodes valence in a distributed fashion, and (2) if the multivariate model does not significantly outperform the univariate model, this would indicate that valence is represented in a univariate fashion, by the global activity level of the amygdala.

As shown in Figure 2A, both the univariate and multivariate models were able to predict valence, with R = 0.157 for the univariate model and R=0.164 for the multivariate model, at p < 0.001 for both models according to a random permutation test. This result indicates that valence information is indeed encoded in the activity of the amygdala. However, a comparison of performance between the multivariate regression model and the univariate regression model revealed no significant difference in predictive accuracy (p = 0.42), suggesting that valence encoding in the core amygdala is primarily driven by its global activation levels rather than by the distributed activities.

**Figure 2.**
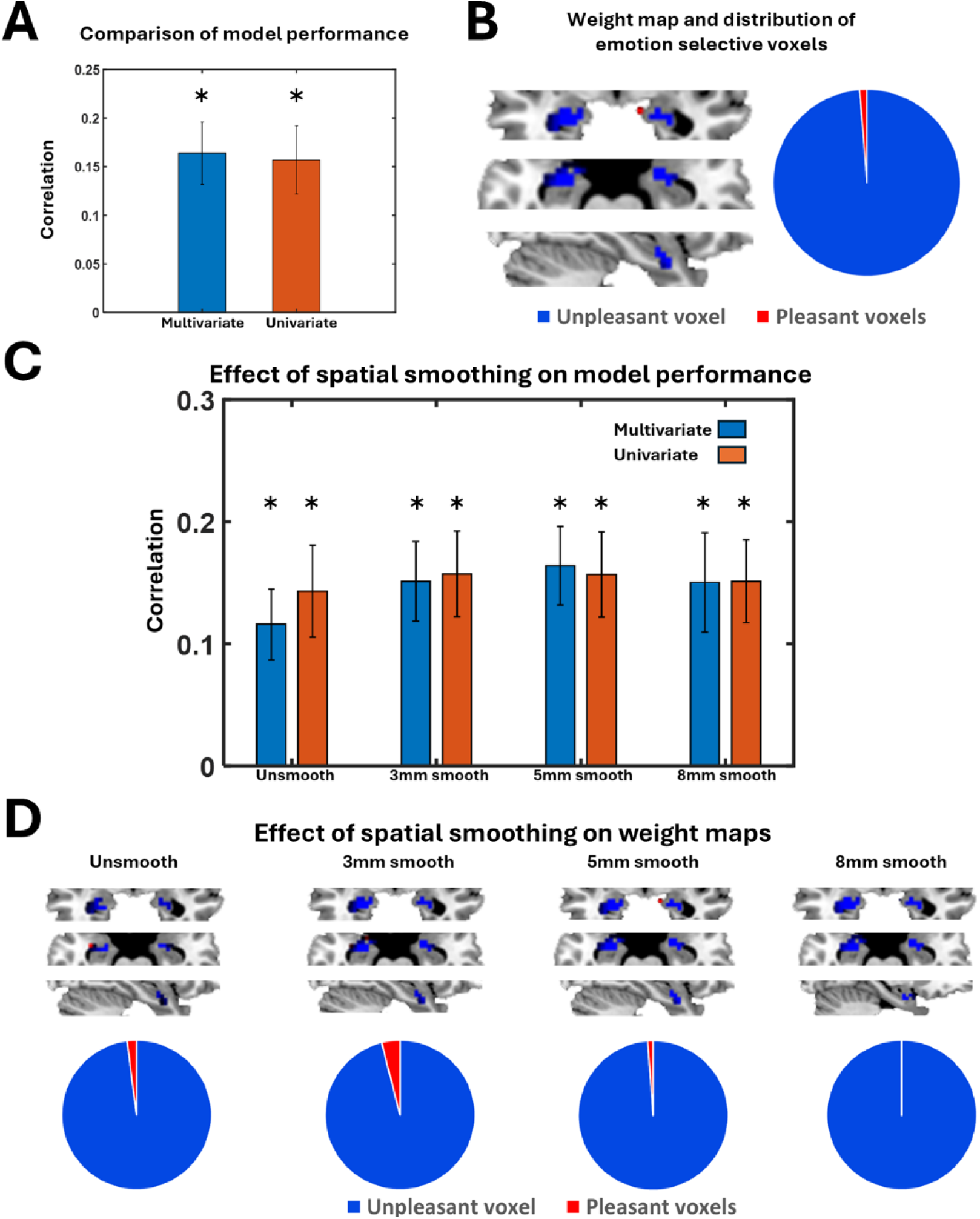
Hypothesis testing in the core amygdala. **(A)** Comparting the performance of the multivariate regression vs the univariate regression models. Both models are able to predict valence (*: p<0.001). There is no significant difference between the two models in valence prediction performance. **(B)** Weight map derived from the multivariate regression model. The voxels are predominantly selective for negative emotions (blue). **(C)** Effect of spatial smoothing on model performance. **(D)** Effect of spatial smoothing on weight maps.

To test this idea further, we investigated the preference of amygdala voxels toward positive or negative emotions by conducting an additional analysis using the weight map derived from the multivariate regression model. The original beta values in the multiple regression model were transformed into a physiologically interpretable voxel map using the method introduced by Haufe et al. (2014). As shown in Figure 2B, 98.8% of core amygdala voxels were found to preferentially respond to negative stimuli, namely, higher BOLD activity for negative stimuli than positive stimuli. In contrast, only 1.2% of voxels preferentially responded to high-valence stimuli, i.e., higher BOLD activity for positive stimuli than negative stimuli. The weight map analysis thus suggests that, unlike single-neuron studies where distinct clusters of positive-selective and negative-selective neuronal ensembles have been observed (Paton et al., 2006), the core amygdala’s representation of valence, spatially resolved at the fMRI voxel level (millimeter in scale), appears to be dominated by voxels selective for negative stimuli. This predominance, which is consistent with the result showing no difference in performance between multivariate vs univariate models, supports the idea that valence in the core amygdala is encoded by its global activity levels rather than by distributed, valence-specific activity patterns.

In multivariate pattern analysis, concerns may arise as to the role played by spatial smoothing, which may influence the detection of distributed patterns. To address this, we applied the same regression analysis to data that underwent varying degrees of spatial smoothing: unsmoothed, and smoothed using 3mm, 5mm, and 8mm full-width half-maximum Gaussian kernels. As shown in Figure 2C, the results consistently demonstrated that the linear regression performance, regardless of the smoothing kernel size, was significantly above chance level. More importantly, there were no statistically significant differences in valence prediction performance between the multivariate and univariate models for all the smoothing levels. Additionally, the proportion of voxels with a negative preference exceeded 95% in weight maps across all smoothing conditions (Figure 2D), indicating that the findings are stable and robust against varying levels of spatial smoothing.

### Expanded amygdala

The core amygdala appears to be relying on a univariate code to represent valence. To explore whether a more spatially distributed valence representation applies to broader definitions of the amygdala, we considered expanded amygdala ROIs using masks generated by lowering the probability threshold for voxel inclusion (i.e., >30%, >10%, and >0%); the most expanded ROI was formed by including >0% voxels plus another layer of voxels beyond the 0% boundary (Figure 3A). The same modeling framework was applied to each of these expanded amygdala ROIs. The results revealed that univariate regression performance gradually declined as the ROI size increased, whereas the performance of multivariate regression showed a slight improvement, and importantly, the difference in model performance reached statistical significance when the amygdala mask was expanded by one voxel beyond the >0% probability boundary (Figure 3B). These results suggest that when the core amygdala is expanded into a larger spatial extent, emotional valence is increasingly encoded in a distributed fashion, such that the population code becomes more applicable. The weight map analysis further revealed that as the amygdala definition became broader, the proportion of voxels selective for pleasant stimuli became larger: from 1.2% for the core amygdala to 18.9% for the most expanded amygdala ROI, and that the voxels selective to pleasant stimuli are predominantly located outside the core amygdala (Figure 3C). This is further demonstrated in Figure 3D, where positively weighted voxels encircle the amygdala’s core, while the core itself is mainly composed of negatively weighted voxels (see Figure 2B).

**Figure 3.**
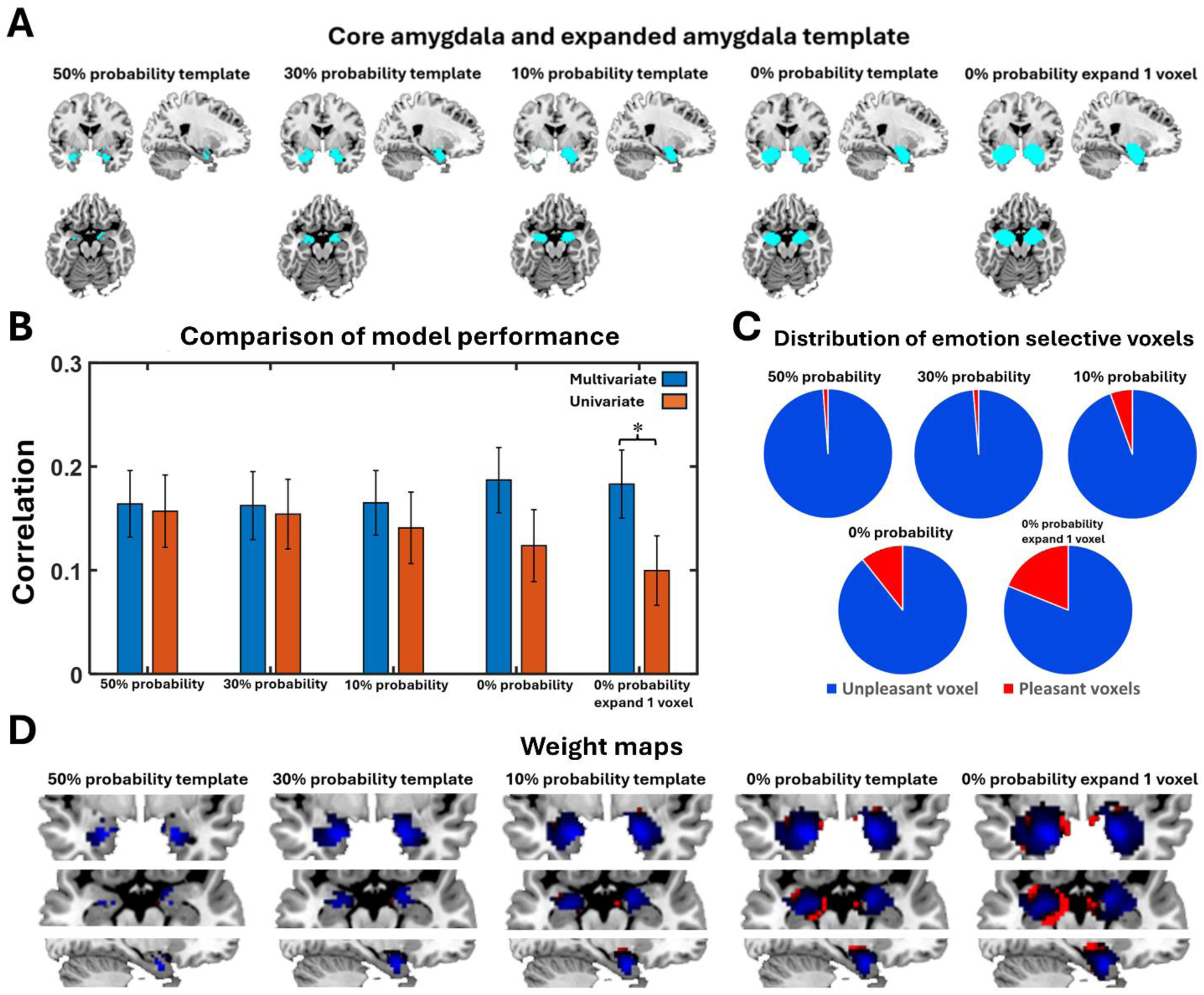
Hypothesis testing in expanded amygdala ROIs. **(A)** Visualization of different expanded amygdala ROIs. Note that the 50% ROI is the same as the core amygdala, which is included here for easy comparison across ROIs. **(B)** Comparison of model performance in predicting valence. **(C)** Distribution of positive-selective and negative-selective voxels. **(D)** Visualization of weight maps for different amygdala ROIs.

### Voxel-level emotion selectivity

Further analysis was carried out to elaborate on the above weight map based results. Whereas linear regression analysis is a commonly applied technique, the weight map analysis is not common applied. Yet, it is the basis for defining stimulus selectivity at the voxel level, which is an integral part of our approach. Although the multivariate model yields weights for voxels (the beta values), these weights, referred to the original weights here, are difficult to interpret as they do not consistently define selectivity; noise and covariation among voxels can render the original weight map uninterpretable (Haufe et a., 2014). For instance, consider a simple two-voxel system. Let voxel 1 and voxel 2 be described by *x₁(n) = s(n) + d(n)* and *x₂(n) = d(n)*, respectively, where *s(n)* is the signal of interest and *d(n)* is shared noise. In this example, voxel 1 contains signal and should be given high weight, and voxel 2 contains just noise and should be given low weight. However, a linear model that best predicts *s(n)* assigns weights [1, –1] for voxel 1 and voxel 2, since the best estimate of the signal is ŝ*(n) = x₁(n) – x₂(n)*. Thus the magnitude of the original weight is high for both voxels. Moreover, the –1 weight for voxel 2 does not indicate true negative selectivity; this high weight merely reflects that voxel 2 contributes to the linear model by canceling the shared noise in voxel 1. The true weight map should instead be [1, 0]. To recover physiologically interpretable selectivity, we used the transformation proposed by Haufe et al., (2014): A = cov(X) × W, where X is the voxels × samples data matrix, W is the original weight vector, and A is the interpretable weight map indicating the weight. In our two-voxel example, assume *s(n) ∼ N(μ = 3, σ² = 1)*, *d(n) ∼ N(μ = 0, σ² = 1)*, The covariance matrix S of [*x₁, x₂*] is : 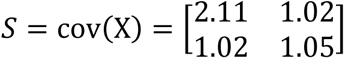. Right-multiplied by W=[1, –1] ^T^ leads to A = [1.09, − 0.03]^T^, which, referred to the adjusted weights, correctly reflects strong positive selectivity of voxel 1 and negligible selectivity of voxel 2 (Cui et al., 2025).

Figure 4A and 4B show the original weight map and the adjusted weight map on the most expanded amygdala ROI (>0% voxels plus one layer of voxels beyond the 0% boundary). For voxels with negative weights (blue), the difference between the BOLD signals that were averaged across unpleasant images and the BOLD signals that were averaged across pleasant images was computed, and for voxels with positive weights (red), the difference between the BOLD signals that were averaged across pleasant images and the BOLD signals that were averaged across unpleasant images was computed. For the original weight map, whereas blue voxels showed a significant BOLD difference (p < 0.0001), suggesting selectivity for negative images, red voxels did not show a significant BOLD difference (p = 0.2564) (Figure 4C). In contrast, for the adjusted weight map, both blue voxels and red voxels showed significant difference in BOLD activation (p < 0.001 for blue voxels, p = 0.0324 for red voxels) (Figure 4D), suggesting negative selectivity for blue voxels and positive selectivity for red voxels. In addition, for the adjusted weight map, we rank-ordered the pictures from 1 to 60 according to the ascending value of the valence, averaged BOLD signals for each picture across blue voxels and across 20 subjects, and plotted the resulting 60 values as a function of the picture index. As shown in Figure 4E, the slope of BOLD activation as a function of increasing valence in negatively weighted voxels was –0.0057 ± 0.0011, which was significantly less than zero (p < 0.0001), indicating increase in activity in blue voxels with more negative stimuli. For positively weighted voxels, the slope was 0.0019 ± 0.00096, which was significantly greater than zero (p = 0.03) (Figure 4F), indicating increase in activity in these voxels with more positive stimuli.

**Figure 4.**
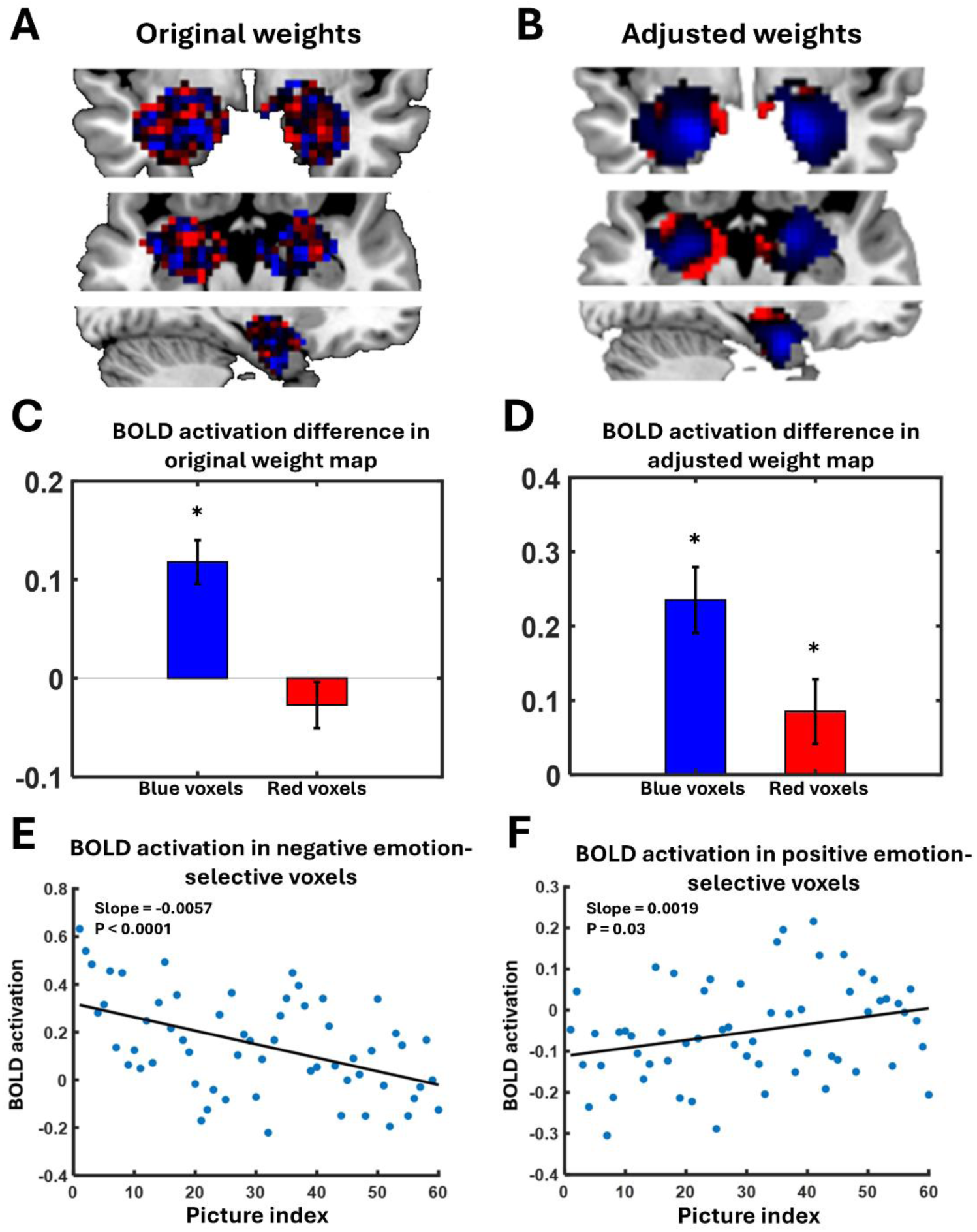
Additional weight map based analyses. **(A)** Visualization of the original weight map on the most expanded amygdala ROI (one layer of voxels beyond the 0% boundary). **(B)** Visualization of the adjusted weight map in the same ROI. **(C)** and **(D)** Selectivity analysis on the original versus the adjusted weight map. Blue: BOLD activation difference between unpleasant and pleasant pictures in blue voxels. Red: BOLD activation difference between pleasant and unpleasant pictures in red voxels. * indicates significantly greater than 0. **(E)** and **(F)** Pictures were rank-ordered from 1 to 60 in ascending valence value (i.e., picture 1 being the most unpleasant and picture 60 being the most pleasant). BOLD activation evoked by each picture as a function of picture index in blue voxels **(E)** and in red voxels **(F**).

## Discussion

The amygdala is a key brain structure for processing emotional information. As valence is a key characteristic of an emotional stimulus, it is important to understand how valence is represented in the amygdala. Two hypotheses have been put forward, based on animal studies, to account for the role of the amygdala in representing the affective valence of sensory input: the univariate representation model and the distributed representation model. To test these hypotheses, we recorded fMRI data during viewing of IAPS pictures and applied both univariate and multivariate methods to multiple anatomical definitions of the amygdala: (1) a widely used amygdala template referred to the “core amygdala” and (2) progressively enlarged versions of the core amygdala (“expanded amygdala”), and reported the following findings: (1) in the core amygdala, both the univariate and the multivariate models equally predicted the normative valence of the IAPS pictures and almost 100% of the core amygdala voxels were selectively coding negative valence and (2) for the most expanded amygdala (>0% probability voxels extended further by one layer of voxels), the multivariate model significantly outperformed the univariate model in predicting the normative valence of the IAPS pictures and the voxels located in the regions surrounding the core amygdala were predominantly selective for positive valence. These results suggest that the neural representations of affective valence transitions from univariate encoding to multivariate encoding as the extent of the amygdala definition increases.

Population coding of valence in the amygdala is well-established in animal models. Electrophysiological recordings from associative-learning paradigms have shown that different ensembles of neurons—particularly within the basolateral nucleus—are selectively tuned to positive (rewarding) versus negative (aversive) stimuli (Belova et al., 2008; Paton et al., 2006). These positive- or negative-selective neuronal ensembles, potentially specified by genetic factors (Kim et al., 2016), project to distinct downstream targets: the nucleus accumbens for reward and the central nucleus of the amygdala for aversion (Smith et al., 2021), and are important components of distinct neural circuits mediating approach versus avoidance behaviors (O’Neill et al., 2018).

Although it is reasonable to expect that these mechanisms are preserved in the human amygdala, they have rarely been empirically tested. Previous human studies have typically focused on overall amygdala activity rather than on patterns of activity (Wang et al., 2005; Schneider et al., 1997). One possible reason is the dominance of the univariate methodology in neuroimaging and the attendant limitations. Over the past two decades, machine learning based MVPA has provided insights on the pattern level encoding of cognitive variables that are not possible with the traditional univariate approaches (Visser et al., 2013; Bo et al., 2021; Bo et al., 2022; Kragel & LaBar, 2014). A critical distinction between multivariate and univariate models is that the multivariate model leverages a weighted combination of voxel-level activations, whereas the univariate model uses voxel-averaged activations, which disregard the differences in patterns of activity. Mathematically, machine learning classifiers can maximize prediction performance by incorporating distinct variance information from multivariate features (e.g., fMRI voxels or EEG channels; see Bo et al., 2021; Bo et al., 2022). In this work, we combined the univariate model and the multivariate model to test the two hypotheses. As shown in Figure 1B, if the univariate model performs similarly in terms of predicting valence as the multivariate models, then the univariate activation hypothesis is being supported because no additional information is contained in the pattern of activation. In contrast, if the multivariate model outperforms the univariate model in predicting valence, then that is taken as evidence to support the population coding model because additional information is being gained by considering the pattern of activation. In the core amygdala, which is considered a conservative definition of the amygdala, we found no significant difference between the performance of the univariate vs the multivariate regression model and showed that most voxels in this region are selective for negative valence. This suggests that, at least at the voxel level spatial resolution, the amygdala encodes valence information in a global up-and-down fashion rather than in a distributed pattern, in agreement with a recent MVPA study that failed to detect distinct neural representations for different affective states in the core amygdala (Varkevisser et al., 2023). As we gradually expanded the spatial extent of the amygdala by including voxels with lower anatomical probability, intriguingly, the performance difference between univariate and multivariate regression analyses increased in favor of the multivariate model and eventually the difference reached statistical significance for the most expanded amygdala ROI, and the voxels that are included in the final expansion process are mainly selective for positive valence. This suggests that as the amygdala definition expands, valence information is increasingly encoded in a distributed fashion rather than in a global up-and-down modulation.

At first blush, our findings seem to be at variance with animal studies that have identified discrete populations of amygdala neurons that are separately tuned to negative or positive stimuli (Kim et al., 2016). Two potential factors may account for the discrepancy between animal and human findings. First, a fMRI voxel contains a large number of neurons, and thus voxel activity reflects a gross summary of more nuanced neural population activities (Ekstrom, 2010; Mukamel et al., 2005). Within voxels that appear selective for unpleasant stimuli, both positive- and negative-selective neural populations may coexist, and it is just that the proportion of positive-selective neurons is likely lower than that of negative-selective neurons; for a related single-neuron study, see (Iwaoki & Nakamura, 2022). Consequently, when we say that almost all the voxels within the core amygdala are selective for negative stimuli, it may mean that the negative-selective neurons are more numerous and their activity is more dominant than the positive-selective neurons. Second, there is accumulating evidence that the core amygdala and its surrounding regions jointly process emotional information (McGaugh 2004; Phelps 2004). Previous meta-analyses have shown that, in nearly half of the reviewed studies, the voxels that showed peak activation to emotional stimuli were actually located outside the core amygdala. Specifically, approximately 11% of these voxels lie in the core region of the hippocampus (Ball et al., 2009), and only 8% of activation peaks in response to positively valenced stimuli were located in the core amygdala (Ball et al., 2009). Our result from the most expanded amygdala revealing that positive-selective voxels were primarily located in regions of the amygdala with low anatomical probability, namely they are located outside the core amygdala, are in broad agreement with this literature. It is likely that the encoding of positive emotions relies more heavily on regions adjacent to, or functionally connected with the core amygdala, rather than in the core amygdala per se.

Neuroimaging studies debate whether cognitive processes are implemented in isolated, localized brain regions (Jonas & Kording, 2017) or require multiple regions working jointly to form a functional network (Brodersen et al., 2012; Vickery et al., 2011). While it remains challenging to observe such large-scale cooperation at the single-cell level, macro-level approaches using fMRI can offer more generalizable insights into the brain’s integrative processes. In terms of emotional processing, our results reflect the possibility that a larger circuit, including regions such as hippocampus, para-hippocampus and nucleus accumbens, cooperate with the amygdala to encode the emotional significance of sensory input. This is not surprising given that previous studies have extensively demonstrated that the hippocampus and parahippocampus store emotional memory by interacting with the amygdala (Aminoff et al., 2013; Phelps, 2004; Smith et al., 2004). The parahippocampus has additionally been shown to provide contextual information to the amygdala during emotion processing (Aminoff et al., 2013). Furthermore, the nucleus accumbens, hippocampus, and amygdala are integral to the dopamine signaling pathway (Alexander et al., 2021), a critical neurotransmitter system implicated in reward-related processes. Taking the network view one step further, in (Varkevisser et al., 2023)’s study, the whole-brain decoding of emotion shows much better performance than amygdala-based decoding of emotion. These findings suggest that human emotion is an extremely complex cognitive process and the amygdala only functions as part of a broader distributed network, which encompass many parts of the brain, to encode emotional valence, possibly through dynamic interactions with the other regions in the broader network.

## Acknowledgements

This work was supported by National Science Foundation grant BCS2318984 and National Institute of Mental Health grants MH112558 and MH125615.

